# Stand age diversity and climate change affect forests’ resilience and stability, although unevenly

**DOI:** 10.1101/2023.07.12.548709

**Authors:** Elia Vangi, Daniela Dalmonech, Elisa Cioccolo, Gina Marano, Leonardo Bianchini, Paulina F. Puchi, Elisa Grieco, Alessandro Cescatti, Andrea Colantoni, Gherardo Chirici, Alessio Collalti

## Abstract

Stand age significantly influences the functioning of forest ecosystems by shaping structural and physiological plant traits, affecting water and carbon budgets. Forest age distribution is determined by the interplay of tree mortality and regeneration, influenced by both natural and anthropogenic disturbances. Thus, human-driven alteration of tree age distribution presents an underexplored avenue for enhancing forest stability and resilience. In our study, we investigated how age distribution impacts the stability and resilience of the forest carbon budget under both current and future climate conditions. We employed a biogeochemical model on three historically managed forest stands, projecting their future as undisturbed systems, i.e., left at their natural evolution with no management interventions. The model, driven by climate data from five Earth System Models under four representative climate scenarios and one baseline scenario, spanned 11 age classes for each stand. Our findings indicate that Net Primary Production (NPP) peaks in the young and middle-aged classes (16- to 50-year-old), aligning with ecological theories, regardless of the climate scenario. Under climate change, the beech forest exhibited an increase in NPP and maintained stability across all age classes, while resilience remained constant with rising atmospheric CO_2_ and temperatures. However, NPP declined under climate change scenarios for the Norway spruce and Scots pine sites. In these coniferous forests, stability and resilience were more influenced. These results underscore the necessity of accounting for age classes and species-specific reactions in evaluating the impacts of climate change on forest stability and resilience. We, therefore, advocate for customized management strategies that enhance the adaptability of forests to changing climatic conditions, taking into account the diverse responses of different species and age groups to climate.

## Introduction

Over the last decade, the European forest policies (Cerullo et al., 2023) have geared towards sustainable and climate-resilient forests to ensure ecosystem functioning under future climate conditions and the sustained provision of services (e.g., regulation, supply, cultural services), including carbon sequestration for forest-based mitigation strategies (Yousefpour & Hanewinkel, 2016; Nabuurs et al., 2018; Churkina et al., 2020; Favero et al., 2021). The contribution of forests to land-based mitigation actions is shaped by external and internal drivers that influence the ecosystem’s net carbon accumulation. This dependency is uncertain under future changing environmental conditions (Yousefpour & Hanewinkel, 2016). Carbon assimilation and wood production are affected by environmental drivers like precipitation and temperature, light availability, atmospheric CO_2_ concentration (CO_2_), and nitrogen deposition (Barford et al., 2001; D’Andrea et al., 2020, 2021; He et al., 2023). In the temperate biome, the effects of increasing temperature on wood production are ambiguous, with studies reporting both positive and negative impacts (Jump et al., 2006; Potočić et al., 2021). As reported in several studies, the positive synergistic effect of elevated atmospheric CO_2_ levels and climate dynamics has likely accelerated European forest growth in recent decades (Pretzsch et al., 2014; He et al., 2023). Other studies, however, question the strength and future persistence of the CO_2_ fertilization effect on the forest carbon sink due to the emergence of other limiting factors, like water and nutrient availability (Nabuurs et al., 2018; Duffy et al., 2021). Climatic models have predicted that the impacts of future climate change will likely alter temperature and precipitation patterns, subsequently influencing atmospheric vapor pressure deficit and soil water availability (IPCC, 2021). These changes could affect the rates of photosynthesis and plant respiration and, ultimately, plant productivity (Reyer et al., 2014; Yuan et al., 2019; He et al., 2023). In addition, significant impacts on forest structural and biological diversity are anticipated (IPCC, 2021). These changes will interact with environmental factors, influencing ecosystem productivity, resilience, and stability. Therefore, evaluating forest ecosystems’ stability is critical for understanding their resistance to environmental pressures and human-induced disturbances. The sensitivity of forests to climate change underscores the importance of comprehensive conservation strategies to protect these fragile ecosystems. Forest stand heterogeneity in landscapes (Bohn and Huth, 2017), variations in leaf area index (LAI, Vilà et al., 2013), and species compositions (Asner et al., 2003) are all factors influencing forest productivity. Climate change is expected to accelerate tree growth, induce earlier maturation, and shorten tree lifespan (Collalti et al., 2019; Brienen et al., 2020), potentially affecting forests of various age structures differently (Anderson-Teixeira et al., 2013; Colangelo et al., 2021).

The traditional view of forest dynamics in undisturbed ecosystems describes an initial stepwise increase in productivity, followed by stabilization and a slight decline (Kira and Shidei, 1967; Odum, 1969). While this view has been found robust under current climate conditions, this concept has been scarcely re-evaluated under climate change scenarios. Notably, future environmental shifts (e.g., climate, nutrients, atmospheric CO_2_ levels) might impact forest stands, or ‘cohorts’ of different ages differently, as both gross primary production (GPP) and plant respiration —key components of the autotrophic carbon budget— vary with stand development (Drake et al., 2011; Goulden et al., 2011; Collalti et al., 2020a). The net balance between photosynthesis and respiration, i.e. net primary production (NPP), being climate- and age-dependent, is likely to respond differently across age spectra under future climatic conditions because these two processes respond differently (Goulden et al., 2011; Collalti 2020a, 2020b).

Given the significant role of silvicultural practices in shaping the current age distribution of managed (Wulder et al., 2009; Latterini et al., 2023), understanding the tree-level age-sensitivity to climate change in the forest carbon budget is fundamental for developing future climate-smart forest management schemes. A deep understanding of the processes affecting the age dependency of CO_2_ uptake and wood production is essential for devising strategies to enhance forest resilience and stability under climate change.

Despite the potential impact of climate change on forests of all ages, there is a lack of prospective understanding of its interaction with forest age. Unfortunately, only a few models have examined age and the ecosystem carbon budget interplay in a climate change context (Anderson-Teixeira et al., 2013). Process-based models (PBMs) enable investigations into the effects of climate change and atmospheric CO_2_ on different age cohorts within the same geographic location, a task challenging with field measurements. Using a validated biogeochemical, biophysical PBM, this study aims to quantify controls on carbon assimilation in undisturbed forests, considering the effects of climate change and stand age. We also evaluated the sensitivity of NPP, a crucial flux representing the annual net carbon sequestration capacity, to key variables such as mean annual temperature (MAT), atmospheric CO_2_ concentration, and vapor pressure deficit (VPD) across different stand ages under five climate scenarios, including a no-change climate scenario.

The specific objectives are: (i) to explore the direct impact of climate change on forest productivity across various stands, species, and age cohorts in different European regions; (ii) to investigate how forest age might modulate dynamics in response to future climate change; (iii) to assess whether age diversity influences the stability and resilience of future forests under changing climatic conditions.

## Materials and Methods

### Study sites and virtual stands

The study was conducted in three even-aged, previously managed European forest stands: i) the Boreal Scots pine (*Pinus sylvestris* L.) forest of Hyytiälä, Finland (FI-Hyy); ii) the wet temperate continental Norway spruce (*Picea abies* (L.) H. Karst) forest of Bílý Krìz in the Czech Republic (CZ-BK1); and iii) the temperate oceanic European beech (*Fagus sylvatica* L.) forest of Sorø, Denmark (DK-Sor). For each site, daily bias-adjusted downscaled climate data from five Earth System Models (i.e., HadGEM2-ES, IPSL-CM5A-LR, MIROC-ESM-CHEM, GFDL-ESM2M, and NorESM1-M) driven by four Representative Concentration Pathways, namely RCP 2.6, 4.5, 6.0, and 8.5 were available. For more detailed information on the study sites characteristics and climate data, see Collalti et al. (2018, 2019), Reyer et al. (2020), Mahnken et al. (2022), and Dalmonech et al. (2022). The chosen sites have been selected due to their long monitoring history and the availability of a wide range of data sources for both carbon fluxes and biometric data for model evaluation, as well as bias-corrected climate scenarios for simulations under climate change scenarios. In addition, these stands: i) cover a wide climatic gradient, ii) their current state is the result of the legacy of past forest management, iii) they are mainly mono-specific and therefore represent interesting «living labs» to study the effects of climate change on single-species and their productivity, reducing confounding effects which otherwise make models struggle to predict forest growth and carbon dynamics (e.g., Vacchiano et al., 2012; Marèchaux et al., 2021), and they have already been in-depth investigated in the context of climate-smart-forestry silvicultural scenarios (Dalmonech et al., 2022).

The model was forced with the modeled climate under different emission scenarios, corresponding to the RCP atmospheric CO_2_ concentration values for the period 1997 to 2100, ranging from 421.4 µmol mol^-1^ in the “best-case scenario” (RCP 2.6) to 926.6 µmol mol^-1^ of the ‘worst-case scenario’ (RCP 8.5). For comparison purposes, the forest model was forced with a detrended and repeated meteorology and atmospheric CO_2_ concentration from 1996-2006, i.e., the current climate scenario (CCS), considered the current climate’s baseline.

At the beginning of the simulations, we created a Composite Forest Matrix (CFM) following the approach described in Dalmonech et al. (2022) to simulate the potential effect of climate stressors on stands of different ages. The 3D-CMCC-FEM v.5.6 has been run at each site to cover the rotation period of each species (from 1997 to 2099) amid the current climate scenario (fixed atmospheric CO_2_ concentration at the year 2000 of 368.8 μmol mol^−1^) consisting of detrended and repeated cycles of the present-day observed meteorology from 1996 to 2006 and the BAU management practices observed at the site. Data required to re-initialize the model at every tenth of the rotation length were retrieved from each simulation. Hence, ten additional stands were chosen for each age in the composite matrix and added to the CFM. This collection of virtual forest stands was used to set different starting ages at the present day (age_t0_) due, ideally, to the past silvicultural practice and climate. These new stands were used to assess the impact of climate forcing on a full range of age_t0_, as the model has already been shown to be sensitive to forest stand development and the relative standing biomass (Collalti et al., 2018, 2019, 2020a).

### The model, model runs, and results evaluation

The 3D-CMCC-FEM (*Three Dimensional - Coupled Model Carbon Cycle - Forest Ecosystem Module*; Collalti et al., 2014, 2016, 2017; Marconi et al., 2017; Mahnken et al., 2022; Dalmonech et al., 2022, 2024; Testolin et al., 2023) was initialized with the structural attributes of the newly created stands from 1997, which was consequently the starting year of all simulations. Modeled climate change simulations under different RCP-emissions scenarios started to differentiate in 2006 (up to 2100). The simulation runs from the different stand initial conditions, corresponding to different age_t0_ classes, were carried out without management as we are interested in the direct climate impact on undisturbed forest stand response, avoiding the confounding effects of forest management on the responses (for forest management effects, see Dalmonech et al., 2022). A total of 825 different simulations were performed as they are the combination of 5 ESM*5 RCP (4 RCPs + 1 present-day climate scenario) * 11 age_t0_ classes * 3 sites. Results were reported for NPP as it is considered one of the most representative and fundamental variables in the carbon cycle. The 3D-CMCC-FEM model underwent initial assessment against observed climate conditions based on field data availability for GPP and NPP (gC m^−2^ year^−1^) as well as the diameter at breast height (DBH) following the methodology reported by Dalmonech et al. (2022; see “Model validation” in Supplementary Material). We compared GPP and NPP against eddy covariance estimates and ancillary data for the years 1997-2005 for DK-Sor and FI-Hyy and 2000-2005 for CZ-BK1. We also compared the diameter at breast height (DBH) in all sites with field measures.

### Sensitivity, resilience, and stability analysis

To assess the impact of climate drivers on the NPP of different age_t0_ classes, i.e., initial conditions, we performed a sensitivity analysis under all climate change scenarios considered. In this sense, sensitivity refers to the extent to which a system is either negatively or positively impacted. We tested the dependence of NPP on MAT, atmospheric CO_2_ concentrations, and VPD. These variables constitute three pivotal environmental factors that influence crucial growth processes, including the rates of photosynthesis, carbon and water use efficiency, and stomatal regulation. We calculate the slope associated with each forcing variable by fitting a multivariate linear model to account for multi-collinearity among climate drivers. We use NPP as the dependent variable and MAT, atmospheric CO_2_, and VPD as independent variables. The higher the slope in absolute terms, the higher the sensitivity to that forcing variable.

Theoretical research has shown that when systems approach a point of no return, where a significant and irreversible change is likely to occur, they become less capable of handling disruptions and preventing significant shifts in their behavior (Zampieri et al., 2021). This capacity to deal with perturbations is often called resilience. It has been suggested that this reduced ability to bounce back can be observed through the increased temporal autocorrelation (TAC) in the system’s behavior over time (Scheffer et al., 2009; Forzieri et al., 2022), reflecting a slower recovery process caused by critical changes at these tipping points, a process also called Critical Slowing Down (CSD)(Smith & Boers, 2023). Another suggested predictor of approaching a tipping point is the temporal variability (Carpenter and Brock, 2006; Smith & Boers, 2023), which can be expressed as the variance or standard deviation of the state variable (i.e., NPP) along time. An increase in temporal variance can foreshadow a regime shift and a loss in stability.

In this study, we calculated the long-term resilience and stability for each scenario and age_t0_ as the 1-lag TAC and standard deviation (SD) of the whole time series, respectively. We also computed the 1-lag TAC and standard deviation of detrended NPP time series within a 20-year moving window for each age_t0_ and scenario to assess the interaction between resilience and stability during the simulation period.

## Results

### Effect of age_t0_ classes and climate change on productivity

At the CZ-BK1, NPP values vary under CCS from ∼350 to ∼225 gC m^−2^ year^−1^, while from ∼370 to ∼210 gC m^−2^ year^−1^ similarly across all RCPs (Figure 1A). Also, at CZ-BK1, NPP peaks in the younger age_t0_ classes, independently of the climate scenarios considered (i.e., 12- to 36-year-old class). At CZ-BK1, any of the combinations of age_t0_ x RCPs shows a reduction of NPP compared to CCS. At FI-Hyy, younger age_t0_ classes started with very low NPP values (∼120 gC m^−2^ year^−1^), step wisely increasing and then stabilizing at the middle of the century at ∼420 gC m^−2^ year^−1^ and then declining, more in the oldest age_t0_ classes, from ∼250 gC m^−2^ year^−1^ under RCP 2.6 to ∼230 gC m^−2^ year^−1^ under RCP 8.5 (Figure 1B). At FI-Hyy, the more productive cohorts are clustered around the first third of the age_t0_ classes (i.e., 28- to 56-year-old). Similar to CZ-BK1, at FI-Hyy NPP shows a reduction under RCPs when compared to CCS, except for RCP 6.0 (see Table S3 in Supplementary Material). Last, at DK-Sor, results show different patterns concerning other sites, with NPP at the end of the century varying from ∼800 gC m^−2^ year^−1^ (under RCP 8.5) to ∼560 gC m^−2^ year^−1^ (under CCS) and then smoothly increase toward the end of the century as the severity of the scenario increase in all age_t0_ classes (Figure 1C). Mean annual NPP has the maximum values for the younger age_t0_ classes, with the peak for the 14-year-old. Different from the other sites, at DK-Sor, in all age_t0_ classes, higher RCPs show proportionally increased NPP compared to the CCS scenario (see Table S3 in Supplementary Material).

**Figure 1.**
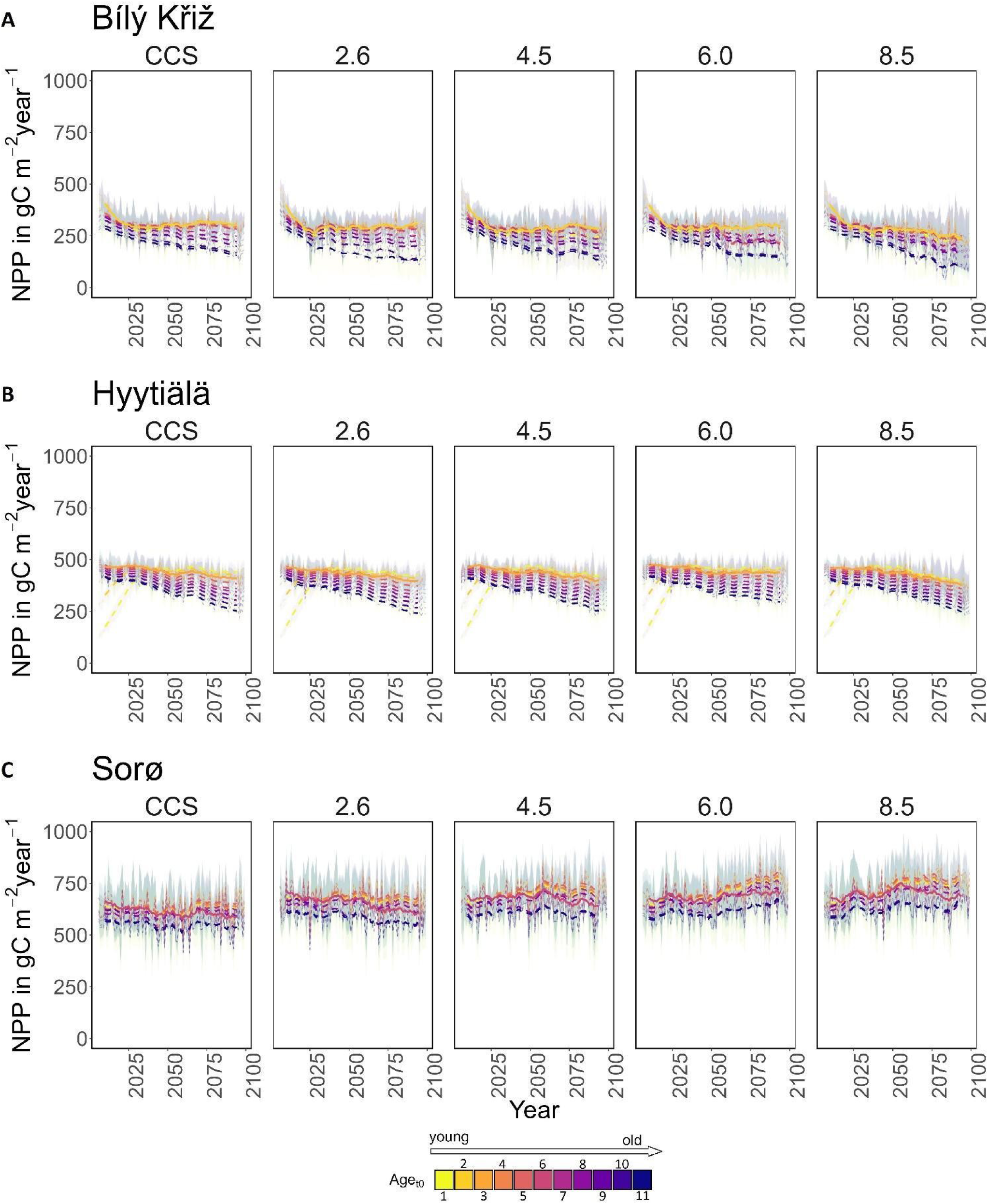
NPP (gC m^−2^ year^−1^) for age classes at the three sites in all scenarios along the simulation period (2006-2099). Lines represent the moving average of 10 years. The solid line corresponds to the real stand, while the dotted line to the virtual one. The shaded area represents two standard deviations from the mean predictions of the five ESMs models. In the legend– for CZ-BK1, FI-Hyy and DK-Sor, respectively – the age_t0_ classes represented by 1) are 12, 14, 14 years-old classes and so on until 11) are 120, 140, 140 years-old classes.

Overall, in all sites and climate conditions, the model effectively captures the typical age-dependent patterns of NPP with relatively stable productivity at young and intermediate ages and a more pronounced decline in older forests. The two conifer stands show a sharper growth decline toward the end of the century across all climate scenarios. This trend was not observed in broadleaf stands.

### Climate sensitivity

The results of the sensitivity analysis are shown in Figure 2 for each site and the three forcing variables considered (MAT, atmospheric CO_2_ concentration, and VPD). At the CZ-BK1 site, under all climate change scenarios, the relationship between productivity and temperature MAT was always negative, with the least sensitive age_t0_ classes being the oldest ones (108-120 years old) in all scenarios except for RCP 2.6, where all classes exhibit a similar sensitivity to MAT. On the contrary, Norway spruce showed a positive relationship with the atmospheric CO_2_ except under RCP 2.6 (i.e. the less enriched CO_2_ scenario), and the least sensitive age_t0_ classes were the middle ones (60-86 years old). Furthermore, as the climate scenario intensified, Norway spruce exhibited an increased sensitivity to elevated temperature and CO_2_ levels. Similar to MAT, the relationship between NPP and VPD was negative under all scenarios for most age_t0_ classes (only the youngest classes showed a positive relationship with VPD under RCP 4.5-8.5). However, the sensitivity increased almost linearly with age_t0_.

**Figure 2.**
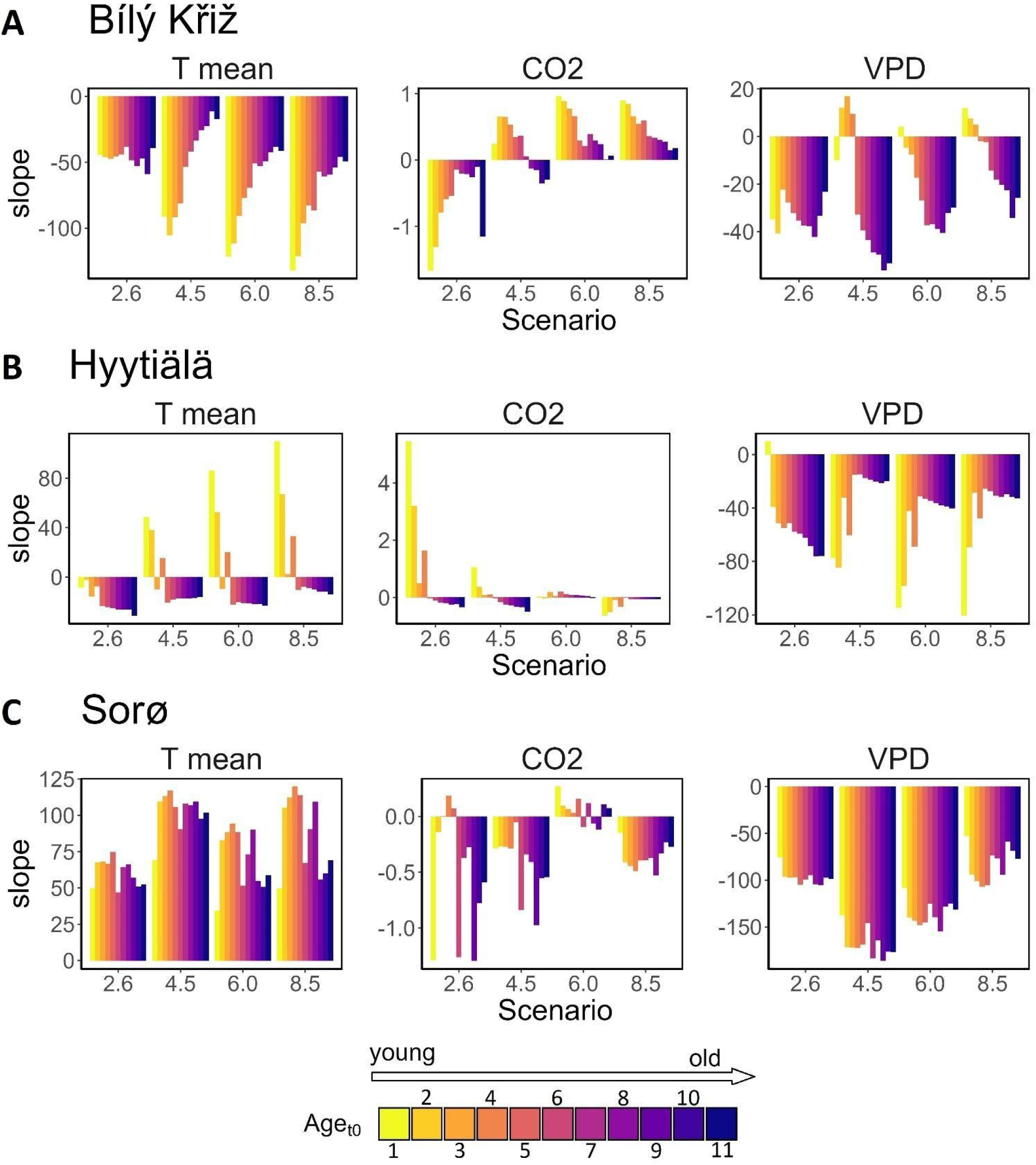
Sensitivity of NPP to three forcing variables (mean temperature, atmospheric CO_2_ concentration, and vapor pressure deficit) in terms of the slope of the regression line resulting from a multivariate linear model. In the legend– for CZ-BK1, FI-Hyy and DK-Sor, respectively – the age_t0_ classes represented by 1) are 12, 14, 14 years old classes and so on until 11) are 120, 140, 140 years-old classes.

Scot pine at FI-Hyy showed similar behavior to Norway spruce but with opposite trends, exhibiting a positive relationship between NPP and MAT only for the youngest age_t0_ (14-28 years-old). The same holds for the atmospheric CO_2_ concentration, but the sensitivity decreases with the severity of the scenario. Similarly to Norway spruce, the sensitivity to VPD generally increases with the age_t0_ but with the highest value for the youngest classes.

At DK-Sor, the beech forest exhibits different patterns with respect to the other sites, featuring a consistently positive correlation between NPP and MAT (but with no clear patterns across age_t0_ and scenarios). On the contrary, a negative relationship was observed against the atmospheric CO_2_ for most age_t0_ classes under all climate change scenarios, except for RCP 6.0 (where the beech exhibited the least sensitivity). The sensitivity to VPD did not exhibit a clear pattern when comparing different age_t0_. However, the highest sensitivity was reached under RCP 4.5.

The findings from the sensitivity analysis indicate that younger age_t0_ classes in the coniferous forests are more vulnerable to variations in climate variables and atmospheric CO_2_ concentration compared to beech forests. Specifically, NPP in conifers demonstrates an inverse correlation with elevated temperature and VPD, whereas in beech forests, the relationship is characterized by a direct proportionality. Overall, NPP sensitivity to forcing variables mildly increases with the severity of climate change in coniferous stands, while the beech forest reaches the highest sensitivity (in particular to MAT and VPD) under RCP 4.5.

### Resilience and stability

The resilience and stability analysis results for each site are reported in Figures 3 and 4, respectively. Figure 5 shows the temporal variation of stability and resilience calculated within a moving window of 20 years. At the CZ-BK1 site, Norway spruce shows increased NPP resilience and stability, progressing from younger to older stands. This observed trend is consistent across all climate scenarios, but as the severity of the scenario increases, resilience tends to increase while stability decreases. In CZ-BK1, younger stands exhibited an initial phase characterized by higher instability and lower resilience, converging over time to conditions comparable to those observed in older stands.

**Figure 3.**
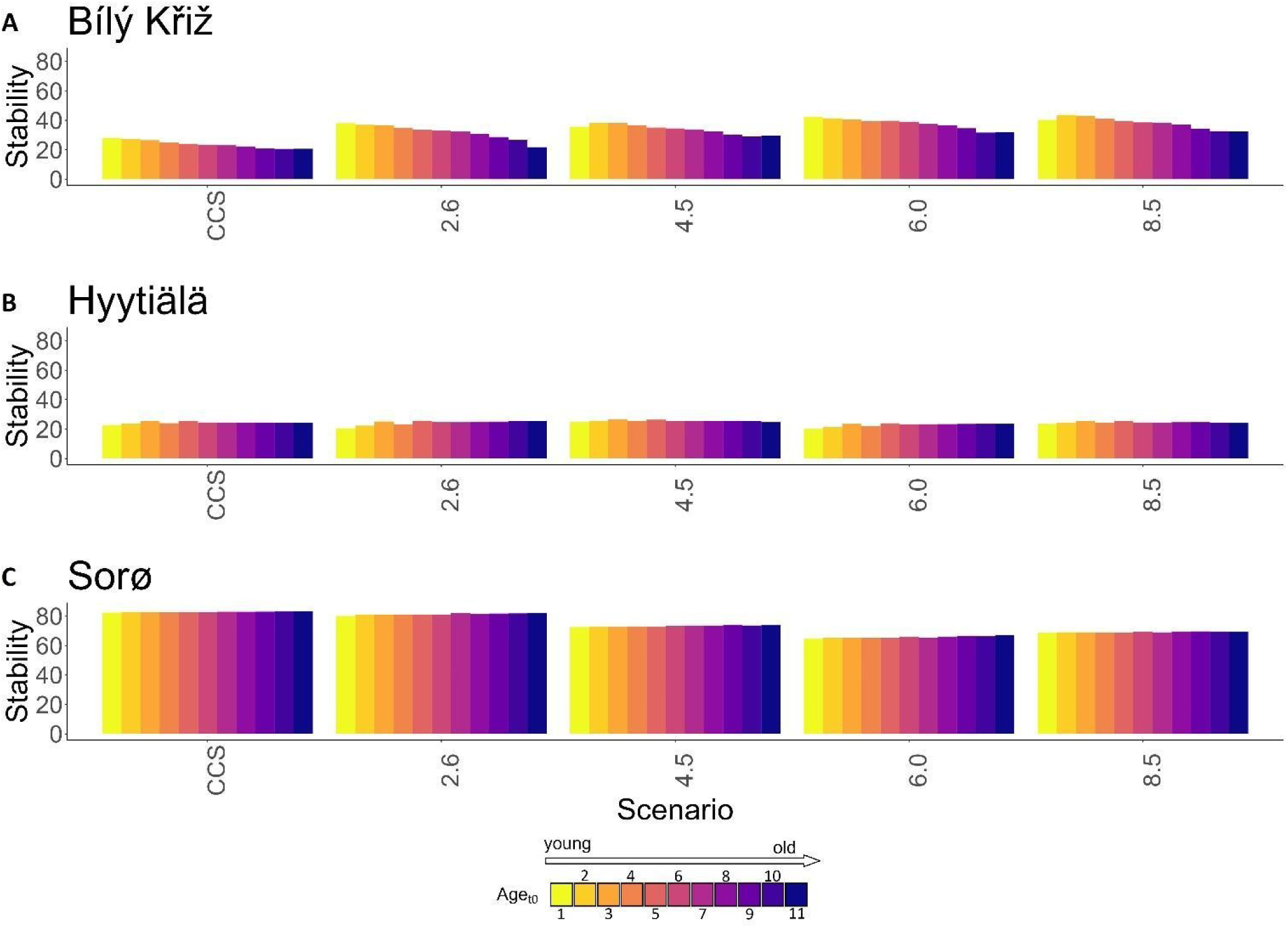
stability (standard deviation of NPP) of each age_t0_ class at the three sites in the five climate scenarios. In the legend– for CZ-BK1, FI-Hyy and DK-Sor, respectively – the age_t0_ classes represented by 1) are 12, 14, 14 years-old classes and so on until 11) are 120, 140, 140 years-old classes.

**Figure 4.**
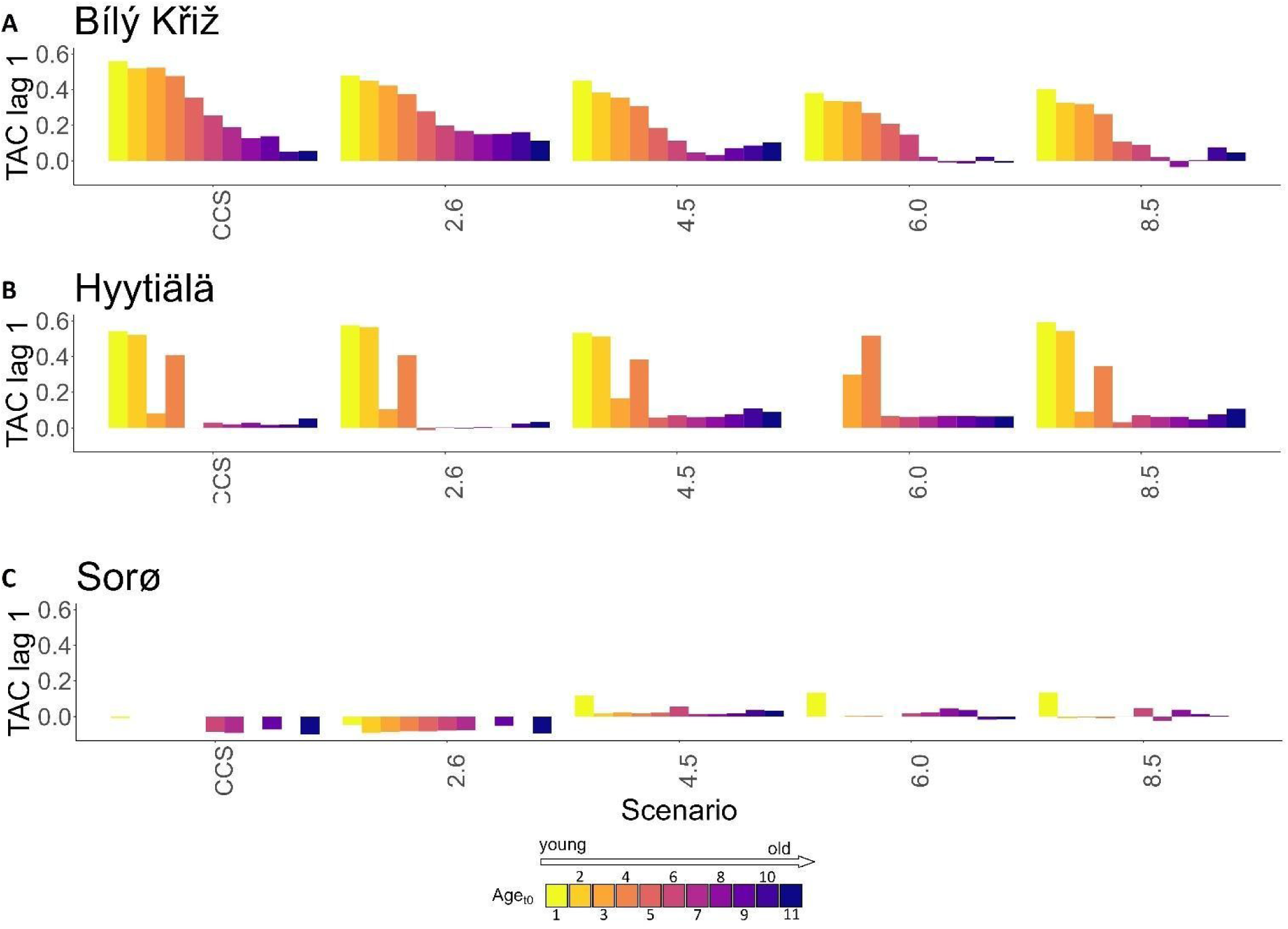
resilience (1-lag TAC of NPP) of each age_t0_ class at the three sites in the five climate scenarios. In the legend– for CZ-BK1, FI-Hyy and DK-Sor, respectively – the age_t0_ classes represented by 1) are 12, 14, 14 years-old classes and so on until 11) are 120, 140, 140 years-old classes.

**Figure 5.**
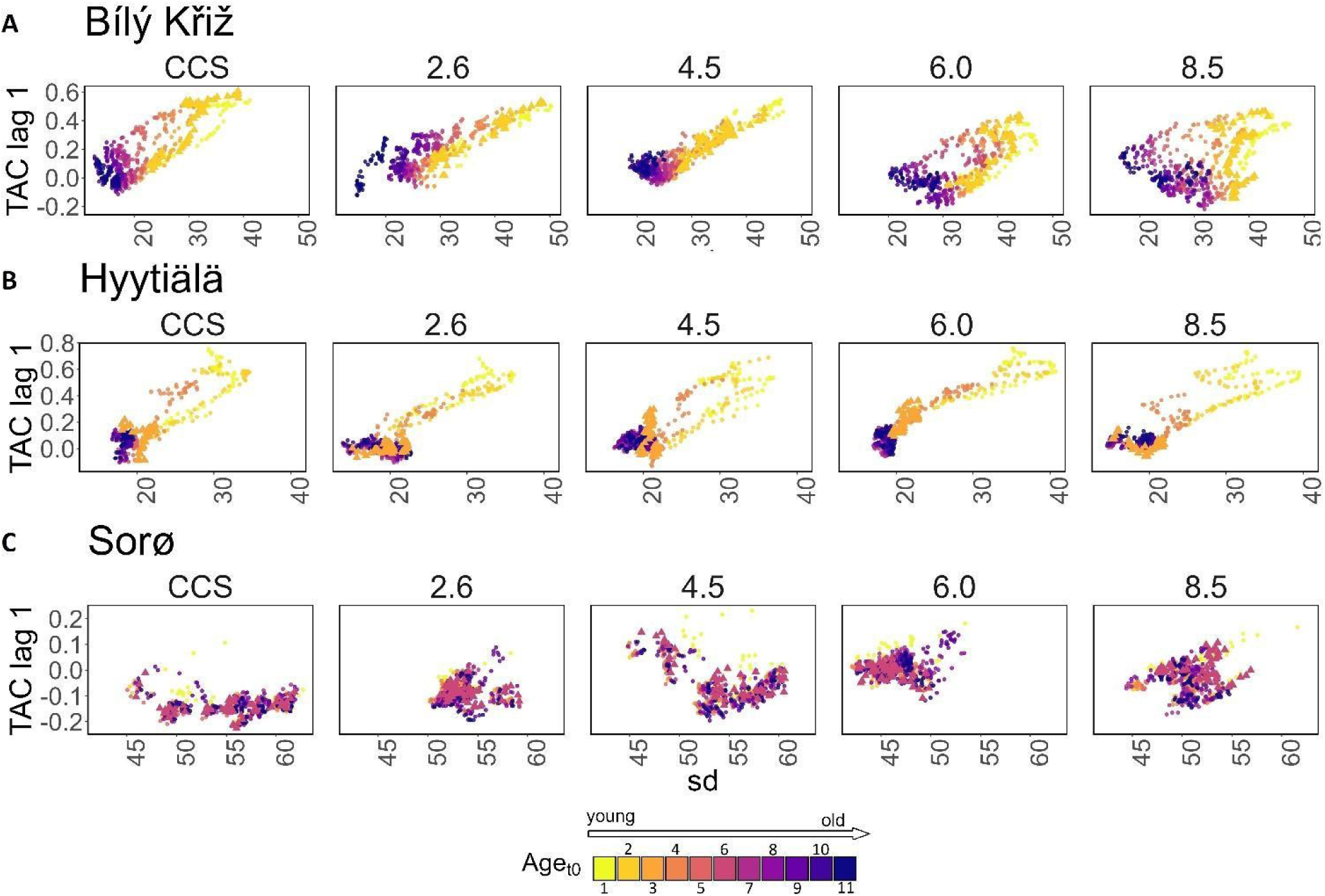
Scatterplot of stability (x-axis) vs. resilience (y-axis) in terms of standard deviation and 1-lag TAC of NPP, respectively. In the legend– for CZ-BK1, FI-Hyy and DK-Sor, respectively – the age_t0_ classes represented by 1) are 12, 14, 14 years-old classes and so on until 11) are 120, 140, 140 years-old classes.

In contrast, older stands demonstrated minimal stability and resilience metrics fluctuations throughout the simulation. The Scot pine at FI-Hyy exhibited a more stable trend in stability, unaffected by the different climate scenarios, with the youngest age_t0_ class showing slightly more stability, although the difference is minimal. Conversely, the younger classes demonstrated the highest long-term autocorrelation, regardless of the climate scenario. As for Norway spruce, the Scots pine showed a similar trend in the temporal variation of stability and resilience. In this pattern, the older classes exhibited remarkable stability and resilience during the simulation period and the tightening of climate scenarios.

Meanwhile, age_t0_ younger than 60 years reach a stable and resilient state at the end of the simulation period. The beech stands at DK-Sor showed an opposite trend concerning the coniferous strands, showing low stability that slightly but steadily increases with the escalation of climate scenarios and high resilience, with minimum differences among age_t0_ classes and scenarios. Throughout the simulation, slight variation in stability and resilience was observed, with only minor variations among the different age_t0_ classes. The results indicate that the most ‘favorable’ climate scenarios were RCP 6.0, characterized by higher stability, and RCP 8.5, in which resilience is highest.

## Discussions

### Stand age and climate effects on productivity

The emerging pattern from the modeling simulation framework shows that regardless of the climate scenarios and in the absence of human intervention, productivity in coniferous stands will decline in the foreseeable future, more likely in the older age_t0_ classes. On the contrary, the future productivity of beech forests at DK-Sor will be expected to increase under most RCP scenarios. These results align with current observations indicating that conifers are suffering from a climate-related decline in NPP (likely due to the greater hydraulic limitation and, therefore, higher sensitivity to the increased soil and atmospheric aridity)(Krejza et al., 2021). The decline was pronounced in conifers compared to broadleaf, especially beech forests, in the most northern species distribution (Del Castillo et al., 2022). Recent studies confirm our results, finding lower relative increment gains for the conifer species Norway spruce and Scots pine (18 and 35%) compared to the deciduous species as European beech and sessile/common oak (75 and 46%)(Pretzsch et al., 2023). Many model simulations have failed to replicate this phenomenon. For example, Reyer (2015) observed that 87% of the simulations run with changing climate and CO_2_ concentration (757 out of 870 simulations) performed with 55 different process-based stand-scale models in Europe predict positive responses in NPP under climate change scenarios and increasing atmospheric CO_2_ concentrations. Reyer (2015) found that only 13% of the simulations show decreasing forest productivity and declining biomass carbon pools, which is opposite to our study. However, in many of the models investigated, as also in other studies considered (e.g., Vitale et al., 2003; Jansson et al., 2008), factors such as photosynthetic acclimation (saturation) to increasing atmospheric CO_2_ concentration, acclimation to rising temperature in autotrophic respiration and drought were not fully and always accounted for, and, often, NPP is considered a fixed fraction of the GPP (see Collalti & Prentice, 2019) or empirically constrained based on current climate conditions (e.g., de Wergifosse et al., 2022). The main consequence of considering NPP as a fixed fraction of GPP instead of prognostically simulating it under climate change scenarios is the increases in NPP *via* increasing GPP, thanks to the CO_2_ fertilization effect (Keenan et al., 2023) and the lengthening of the growing season (Peano et al., 2019). On the other hand, the negative feedback on NPP due to increased autotrophic respiration (for maintenance respiration at tree and stand level) is not considered. As other researchers found (e.g., Musavi et al., 2017), the evidence that the main fluxes of the carbon cycle (i.e., GPP and NPP) are mainly controlled by age suggests that the effects of climate change on forest cohorts could be generally less significant than the effect of age, in particular in coniferous forests, where climate change affects evenly all age_t0_ investigated in this study. These findings align with Foster et al. (2016), who stated in their study that growth variation (in biomass) could be explained mainly by age and size rather than climate variables. In our case, the influence of age_t0_ (that is, the age at the start of the simulation but that increases its effects throughout the model runs) is not a standalone “effect” but rather the result of multiple age-dependent factors. These factors encompass the size of living biomass pools, forest structural traits (e.g., canopy closure, changes in LAI, and alterations in the sapwood volume and mass per unit leaf area), reduction in stomatal conductance (and its effect on photosynthesis), and changes in Non-Structural Carbon usage and allocation. This carbon pool buffers any asynchrony between assimilation, allocation, and plant respiration induced by environmental variability (Collalti et al., 2016, 2020a; Guillemot et al., 2017), thus leading to higher resilience in times of stress (Richardson et al., 2013). The reduction in NPP becomes more prominent with forest aging. This can be caused by an increased maintenance respiration costs due to higher substrate amount and higher enzymatic activity with rising temperatures, although all these mechanisms are yet to be understood (see Collalti et al., 2020a and 2020b). Productivity peaked in the young-intermediate age classes, as also found by Goulden et al. (2011), while in older stands, NPP was drastically reduced. However, it appears that the decline in NPP is not necessarily attributed to the decrease in GPP (thus excluding the fact that NPP is a fixed fraction of GPP), as some suggest (see Drake et al., 2011). Instead, the reduction in NPP is associated with increased autotrophic respiration rate (see DeLucia et al., 2007; Goulden et al., 2011; Drake et al., 2011; Collalti & Prentice, 2019; Collati et al., 2020b) driven by the high maintenance costs for the living, and thus respiring, biomass as identified in this study. The underlying hypothesis is that photosynthesis and dry matter production increase as leaf area increases until the canopy is completely closed. After that, an indirect decrease in soil nutrient availability (Goulden et al., 2011; but not accounted for here), water resources, and an increase in stomatal limitation led to reduced photosynthetic rates; thus, carbon allocation shifts from leaves and roots to aboveground wood production (Guillemot et al., 2017; Meng et al., 2023) but at the expense of increasing respiratory costs. At the stand-level, such a hypothesis is also strengthened by the age-related reductions in population tree density.

### Age and climate-dependent effects of climate change on forest resilience and stability

A growing number of studies have used temporal variance and 1-lag TAC as indicators of CSD (Feng et al., 2021; Smith & Boers, 2023; Forzieri et al., 2022), also referred to as early-warning signals, primarily through remote sensing-based vegetation indices. However, long-term monitoring and prediction of CSD are challenging and not wholly reliable based on satellite time series data. Firstly, remote sensing-based vegetation indices are constrained to the past-present temporal window, are time-delayed, and are often unavailable as suitable long time series. Secondly, remote sensing time series data are often constructed by aggregating multiple data sources with different spatiotemporal and spectral characteristics. This could lead to spurious variations in CSD being misinterpreted as changes in system resilience (Smith & Boers, 2023). Also, satellite data does not yet have a standardized and commonly accepted pre-processing routine to monitor resilience (Smith & Boers, 2023). Finally, in high biomass density regions, optical sensors have a well-known tendency to saturate (Giannetti et al., 2022; Vangi et al., 2021; Smith & Boers, 2023). On the contrary, PBMs can provide long and consistent time series of different key state variables (such as GPP, NPP, carbon stocks, and other structural variables), which can also be extended into the future by simulating the responses of the forest ecosystem to changing environmental conditions, under the most accepted scenarios of climate change. Furthermore, PBMs can be applied to a wide variety of forest ecosystems, assessing the mechanisms of resilience and stability not only in the long-term time dimension but also from the local to continental spatial spectrum.

At the same time, however, the effects of climate change on the carbon budget are highly uncertain and largely debated with models that simulate an overall increase in NPP, possibly due, among other things, to the fixed relation between NPP and GPP (e.g., Reyer 2015) and those models that return an increase in tree and stand respiratory costs over, e.g., the positive effects on GPP because of CO_2_ fertilization and prolonging vegetative seasons (Peano et al., 2019; Keenan et al., 2023). Forest stability and resilience under climate change are even more uncertain.

However, while these two effects, i.e., age and climate change, at different levels, are primarily discussed and reported in the literature, their interaction and effects on forest stability and resilience are not fully investigated. In this regard, it is worth remembering that, as Smith and Boers (2023) pointed out, CSD indicators are not a direct measure of resilience and stability but proxies for state changes. Increasing variance or 1-lag TAC can be caused not only by a resilience loss. This is why it is important to at least investigate the 1-lag TAC and temporal variance together and test whether their behavior is consistent (Ditlevsen & Johnsen, 2010).

In beech stands, the main driver of CSD was the climate. The direct fertilization effect of CO_2_ and the lengthening of the growing season outweigh other limiting factors, which might cause the beech forest to respond more positively to the different climate change scenarios, with productivity increasing up to RCP 8.5 in all age classes. For instance, during the simulation period, under RCP 8.5, there will be an average of 21 more days with MAT>0 °C compared to the CCS scenario. The substantial variability in NPP (as shown in the shaded area of Figure 1C) is emphasized under climate change scenarios in which the growing season of the European beech at this latitude is likely to be short, even shorter than in coniferous stands.

The negative 1-lag TAC under the less warm scenarios (CCS and RCP 2.6) can be caused by the “memory effect”: the whole leaf biomass is replaced at the start of each new growing season, losing the “memory” of the precedent year climate and causing significant differences in productivity from one year to another. Also, the stability was not affected by age at the start of the simulation (age_t0_), while it increased evenly up to RCP 6.0 and then stabilized. The relationship between resilience and stability indicators (e.g., 1-lag TAC and temporal variance, Figure 5C) in the beech stands are not clustered by age_t0,_ suggesting a prominent role of climate in resilience and stability patterns.

Coniferous stands, on the contrary, have a more extended “memory” given to longer leaf turnover rates, preserving the biomass leaf the whole year and for more than one year (about four years for both Scots pine and Norway spruce, Mäkelä et al., 1997 and Pietsch et al., 2005), resulting in a prolonged vegetative phase and less variability in productivity. However, despite this longer “memory” with respect to deciduous species, young needle-leaf stands are generally characterized by a lower LAI (and higher leaf mass per unit leaf area), a shallower root system that brings in less efficient water management, exposing them to drought-induced stress, ultimately causing a loss in stability and resilience (Figure 4A).This evidence is confirmed by the highly clustered resilience-stability scatterplot (Figure 5A-B), suggesting that age_t0_ is the primary driver of resilience and stability in coniferous stands.

In particular, at CZ-BK1, where Norway spruce grows at the edge of its ranges, the effects of environmental changes are amplified, making it the most climate-sensitive species. NPP showed a negative trend under all scenarios, suggesting that the species was already at its optimum and that even slight climatic variations would affect the resistance and resilience of the species. Indeed, several studies highlight the lower stability of the species to water shortage and subsequent drought conditions (Bosela et al., 2021; Mensah et al., 2021; Marchand et al., 2022). Generally, differences among the analyzed species are in line with meta-analyses showing, for instance, how coniferous tend to show greater drought-legacy, which could be mirrored by a higher TAC-1, compared to angiosperms such as beech (see Anderegg et al., 2015)

However, it is interesting that the decrease in stability was compensated by the increase in resilience in the most severe scenarios (RCP 6.0 and 8.5), which could result from dying trees toward the end of the simulation, making perhaps the remaining trees more resilient because of less competition.

Scots pine, despite a similar resilience-stability relationship to Norway spruce, had an intermediate pattern compared to the other two species. The species showed a steady decline in productivity for the older age classes compared to the younger ones, corresponding to lower values of 1-lag TAC (higher resilience), reflecting the instability of young, small trees. Uri et al. (2022), studying the dynamics of carbon storage and fluxes in a Scots pine forest, found that the maximum levels of NPP are observed during the pole and middle-aged developmental stages. Forest stability and resilience are the legacy of past ecosystem states, shaped mainly by species-climate relationships, climate adaptation, and the history of the disturbance regimes (Johnstone et al., 2016). Older forest stands exhibit a higher stability and resilience, probably due to higher robust and stable interaction with climate stressors and better coping with changing environmental factors and conditions than younger stands. Older trees generally have larger carbon pools in sapwood and reserves, which might help buffer the impact of the year-to-year climate variability on tree growth and lead to higher long-term resilience than younger trees (Zweifel et al., 2023). The same argument would hold for beech versus pinus spp, the former characterized by larger carbon pools. In this sense, forest ages could be seen as a “memory” pool of adaptation strategies inherited from past climate and disturbance regimes, dampening the effect of climate change on stability and resilience. Old forests are characterized by species-specific traits perfectly aligned with the climate and disturbance conditions in which they grow, making them more stable and resilient to changes (Reyer et al., 2015).

### Consequences on and for forest management

The diverse behavior exhibited by forests of varying ages, in terms of productivity, stability, and resilience, underscores the potential of utilizing structural heterogeneity at the landscape level to counter climate change impacts while preserving resilience. Heterogeneous and uneven-aged forests profit from increased structural complexity, as evidenced by Jandl et al. (2019), de Wergifosse et al. (2022), and Asbeck et al. (2023). A forest of different age groups integrates young, middle-aged, and mature trees, leading to vertical stratification that augments ecosystem complexity and stability. This stratification fosters gradients in biomass (both deadwood and living aboveground biomass), diverse carbon allocation methods (Merganičová et al., 2019), and ecological niches for flora and fauna, thereby enhancing biodiversity and the capacity to adaptively respond to environmental disturbances (Nolè et al., 2014; Lafond et al., 2014; Pardos et al., 2021).

Our research, intentionally designed to isolate the effects of management strategies, reveals significant implications for climate-adaptive silviculture, even under scenarios where ‘active’ forest management ceases. Functionally, age-diverse forests display a range of physiological and lifecycle traits, enhancing their ability to absorb and store carbon and mitigating the impacts of climate change. Present-day young trees absorb more CO_2_ than their historical counterparts, contributing significantly to carbon capture and efficient biomass production (DeLucia et al., 2007; Campioli et al., 2015; Collalti et al., 2020b), albeit potentially at the expense of a shorter lifespan. Conversely, older trees bolster ecosystem stability and resilience, acting as long-term carbon sinks and regeneration shelters. Thus, maintaining age-diverse forests, facilitated through proactive management, is vital for maximizing the ecological and climatic benefits of structural and functional diversity, balancing each age group’s advantages and drawbacks, as supported by Ehbrecht et al. (2021), Zampieri et al. (2021), and Kauppi et al. (2022). These insights on the interplay between age, climate, resilience, and system stability should be integrated into the functional network approach (Messier et al., 2019), which combines functional traits and landscape connectivity to evaluate functional diversity and its dispersal potential, aiding in recognizing stability and resilience in proactive forest management.

Contrastingly, several studies, including Drake et al. (2011), Jandl et al. (2019), and Dalmonech et al. (2022), argue that unmanaged, aging forests may become less effective against climate change. Forests mainly composed of old-age classes often experience reduced growth and carbon absorption due to competition, nutrient scarcity, and increased disease and pest risks, leading to higher mortality rates. This diminishes their role as carbon sinks (Zhu et al., 2019). Peng et al. (2023) suggest that reducing forest harvesting could be an effective mitigation strategy, though other issues contribute to the debate, such as the ‘higher value’ of short-term mitigation strategies or the unlikely scenario of “no shift” in other emission sources. Arguing for the superior value of near-term emissions and mitigation is questionable, especially considering the short analysis period (2010 – 2050) relative to the lifespan of forest and wood products, which calls for care in interpreting simulation results.

Moreover, the assumption that harvesting cessation will not lead to a shift in emission activities is overly simplistic. Forests have multiple functions, including protection, particularly in Alpine areas. In order to maintain forest functionality and protection against natural hazards (e.g., rockfalls, avalanches, landslides), silvicultural interventions must be applied according to very strict regulations (Moos et al., 2023). Therefore, diversifying forest ages through management not only leverages the rapid growth of younger trees but also balances the benefits and disadvantages of different age classes. A mosaic of varied age patches enhances biodiversity, stability, and resilience to stressors.

Moreover, management prevents forest abandonment, reducing wildfire risks, especially in Mediterranean areas where accumulated combustible material is a concern (Llovet et al., 2009; Ursino & Romano, 2014). Adopting proactive management practices, including selective cutting and promoting natural regeneration and diversity, is crucial. This approach maintains forests of different ages, reflecting varied climate responses, thus ensuring their continued role in climate change mitigation, carbon sequestration, and biodiversity conservation under uncertain climatic futures.

### Limitations

In this study, we did not account for the indirect effects of climate change like altered disturbance regimes and potential extreme events (e.g., fires, insect outbreaks, storms), unless these are already incorporated in the climate scenarios used. This approach intentionally avoids confounding influences on productivity and carbon accumulation estimates. Our focus is on the direct impacts of climate change, specifically increased atmospheric CO_2_ levels, while excluding other environmental drivers such as Nitrogen (N), Phosphorus (P), or Ozone (O_3_). These elements have been recognized as significant influences on forest productivity and vary across different age classes (De Marco et al., 2022a, 2022b). Notably, N deposition rates, which are predicted to double before stabilizing (LeBauer and Treseder, 2008), could enhance fertilization in temperate forests. Additionally, P limitation, rather than N, is more likely to restrict the growth of smaller trees. The increase in N availability might also mitigate the observed decline in NPP in our simulations, as suggested by LeBauer and Treseder (2008). Another significant limitation is the exclusion of heterotrophic and soil responses to avoid additional complexity. This study focuses exclusively on tree-level physiological responses across different age classes, considering the combined effects of altered CO_2_ concentration, temperature, and precipitation. This specific focus is chosen for its direct relevance to forestry management and its potential for short-and long-term mitigation.

Furthermore, we did not include natural regeneration and the possible migration of new species into the study areas. The capacity of tree species change as a climate change mitigation strategy in unmanaged forests might be limited. This is due to the potential discrepancy between species migration or replacement rates and the projected rates of climate change, particularly under more severe RCP scenarios (Settele et al., 2014; Noce et al., 2016).

## Conclusions

In light of growing uncertainty about future climate conditions, forests’ long-term functionality and their consistent delivery of diverse ecosystem services will be unevenly impacted. This impact hinges on several factors: tree species, the geographical position of the forest within the species distribution range, the structural attributes at the stand level, and the stage of stand development. Anticipating climate change, younger forests are expected to experience accelerated growth and earlier maturity but may also have a reduced lifespan. A range of stressors intensified by climate change, including higher temperatures, shifting precipitation patterns, and more frequent extreme weather events, pose significant threats to the stability and resilience of European forests. Grasping the interplay of stability and resilience across different forest age classes is vital for devising strategies to effectively preserve and manage forests in an era of climate change. Adaptive forest management practices that enhance species diversity, support a range of forest ages, and encourage natural regeneration can bolster the resilience of European forests in the face of climate change. Our research underscores the necessity of factoring in both age class and species-specific responses to fully understand the impact of climate change on forest productivity and carbon storage. The varied reactions of tree species to changing climate conditions emphasize the urgent need for customized management and conservation approaches to strengthen forest resilience, stability and adaptability.

## Conflict of Interest

All authors declare no conflict of interest.

## Notes

https://zenodo.org/record/8014127

### Competing Interest Statement

The authors have declared no competing interest.

### Summary of Updates

deleted a typo in the title, no other changes

